# C-Terminal Domain of *Mycobacterium tuberculosis* Glutamate Decarboxylase determines the bacterial stress-adaptive metabolic state that steers macrophage polarisation supporting intracellular persistence

**DOI:** 10.64898/2026.05.05.722917

**Authors:** Anushka Agarwal, Akash Misra, Shalini Saxena, Krishnaveni Mohareer, Anant B. Patel, Sharmistha Banerjee

## Abstract

*Mycobacterium tuberculosis* (*Mtb*) adapts metabolically to survive hostile conditions within alveolar macrophages, but whether specific enzyme-driven metabolic states of *Mtb* link stress adaptation to host immunomodulation remains unclear. Using LC-MRM/MS metabolite profiling and metabolic modeling, we previously identified elevated γ-aminobutyric acid (GABA) production via the GABA shunt during adaptation to oxidative and acidic stress. Here, we show that stress-responsive enzymatic efficiency of glutamate decarboxylase (GadB) in producing GABA, not only mitigates acidic and oxidative stress but also drives macrophage polarisation towards the M2 phenotype, extending its role beyond bioenergetics into host–pathogen cross-talk.

We identify the C-terminal domain (CTD) of mycobacterial GadB as a key regulator. Enzyme kinetics and fluorescence spectroscopy revealed that CTD deletion abolishes pH-dependent regulation of catalytic efficiency (*k*_cat_/*K*_m_) and substrate affinity (*K*_a_). Under oxidative stress, the CTD enhances catalytic efficiency without altering substrate affinity, suggesting condition-specific roles. Accordingly, *in vitro*, *Mtb::Mtb*GadB with higher GABA and ∼40% lower ROS outgrew *Mtb::Mtb*GadBΔCt under acidic and oxidative conditions. These biochemical differences influenced host immunity, wherein infection with high-GABA-producing *Mtb::Mtb*GadB strain induced M2 polarisation, with decreased CD86, reduced RNS, limited endosomal maturation, as indicated by Rab5/Rab7 ratio and Lysotracker-based confocal microscopy, and decreased TNF-α, IL-12p70, and IFN-γ release compared to low-GABA-producing *Mtb::Mtb*GadBΔCt. Consequently, *Mtb::Mtb*GadBΔCt showed higher clearance. We conclude that by enabling pH-dependent activity of GadB, the CTD orchestrates proton consumption, ROS reduction, and GABA-mediated immunomodulation. Distinct structural features of CTDs in human versus *Mtb* GADs highlight the CTD as a selective drug target for pathogen-specific, host-compatible therapies.

**Importance:** This work establishes a direct mechanistic link between metabolic state of *Mycobacterium tuberculosis* (*Mtb*) actively shaping host immune responses, deciding infection outcome. The study identifies the C-terminal domain (CTD) of *Mtb*GadB in steering enzymatic efficiency, regulating intrabacterial GABA production and ROS quenching, determining the stress-responsive metabolic state of the bacteria. Infection with metabolically distinct strains had differential impacts on host immunity, wherein infection with high-GABA-producing *Mtb*::*Mtb*GadB induced M2-polarisation, reduced RNS, limited endosomal maturation, and reduced proinflammatory cytokine release, compared to low-GABA-producing *Mtb::Mtb*GadBΔCt. Consequently, *Mtb::Mtb*GadBΔCt could be cleared, but *Mtb::Mtb*GadB persisted, deciding the fate of the infection. Structural differences between CTDs of human and *Mtb* GADs, make it a therapeutically-exploitable determinant for developing pathogen-specific drugs that remain compatible with the host.

## Introduction

*Mycobacterium tuberculosis* (*Mtb*), the causative agent of tuberculosis (TB), survives for prolonged periods within macrophages despite exposure to multiple host-imposed stresses, including acidic pH, oxidative stress, nutrient limitation, and hypoxia (1–3). Among these, phagosomal acidification and reactive oxygen species (ROS) represent dominant early antimicrobial pressures encountered during infection (4–6). Notably, acidic and oxidative stresses are functionally interconnected, as proton accumulation can influence redox reactions, while oxidative stress can disrupt proton gradients and intracellular pH homeostasis, together imposing a combined metabolic burden on the bacterium (7, 8). These coupled stresses perturb central metabolism and redox balance, necessitating coordinated adaptive responses for the survival of intracellular bacteria (9–11). Consequently, successful persistence of *Mtb* within macrophages relies on metabolic pathways that simultaneously maintain cytosolic pH, redox homeostasis, and energy production.

Metabolic remodeling during infection has emerged as a critical determinant of mycobacterial survival (12). Systems-level studies, including metabolomics and infection models, have revealed stress-induced reprogramming of central carbon metabolism, amino acid utilization, and redox buffering pathways that enable adaptation to hostile intracellular environments (13, 14). Importantly, these metabolic adaptations extend beyond bacterial physiology and intersect with host immune responses, thereby modulating macrophage activation states and inflammatory signaling (15–17). Given that macrophage polarisation is tightly linked to metabolic programs, pathogen-derived metabolites represent a potential mechanism by which *Mtb* may reprogram host immunity to favor its persistence.

Our previous metabolomic analysis identified increased accumulation of γ-aminobutyric acid (GABA) and glutamate in *Mtb* exposed to acidic and oxidative stress conditions (18). GABA is produced from glutamate by the enzyme Glutamate Decarboxylase (GadB), a pyridoxal-5′-phosphate (PLP)–dependent enzyme that catalyzes a proton-consuming decarboxylation reaction (19). In several bacterial systems, this reaction functions as an acid resistance mechanism by consuming intracellular protons and contributing to pH homeostasis (19, 20). In addition, the GABA shunt feeds into central metabolism and can influence redox balance through NAD⁺/NADH-linked reactions (21–24). Recently, various studies have suggested the potential role for GABA in immunomodulation (25–28); however, whether *Mtb*GadB-mediated metabolism can influence host immunometabolic pathways during infection remains an open question. Furthermore, recent studies using non-pathogenic mycobacteria have suggested a role for mycobacterial GadB in intracellular survival (29, 30); however, the biochemical regulation of *Mtb*GadB under host-relevant stress conditions and its potential contribution to host–pathogen interactions remain poorly understood. In particular, the structural determinants that govern enzyme activity, stress responsiveness, and metabolic output have not been defined.

In this study, we investigated the biochemical regulation and functional role of *Mtb*GadB under acidic and oxidative stresses and show that stress-induced catalytic efficiency of *Mtb*GadB aids in adaptation to intracellular stresses and shifts the macrophage immune responses toward an anti-inflammatory state. Importantly, we highlight the significance of the integrity of the C-terminal domain of *Mtb*GadB in intracellular adaptation as well as immunomodulation vis-à-vis enhanced catalytic efficiency. Together, our findings position *Mtb*GadB as a pivotal metabolic node that links bacterial stress adaptation with host immunomodulation. This underscores the essential role of stress-induced bacterial metabolism in mediating metabolic cross-talk between *Mtb* and the host, thereby shaping host–pathogen interactions and revealing the intricate interplay between *Mtb* adaptation and the host immune response.

## Results

### Expression and catalytic efficiency of *Mycobacterium tuberculosis* GadB are enhanced under acidic and oxidative stress conditions

Building on our earlier observation that intrabacterial GABA and glutamate levels rise under *in vitro* infection-mimicking conditions (18), we examined the expression of key enzymes of the GABA shunt pathway in pathogenic *Mtb* strain (*H37Rv)* during acidic and oxidative stress. Mid-log phase cultures were exposed to acidic (pH 5.5) or oxidative (10 mM H₂O₂) conditions. Transcript levels, quantified by qRT-PCR, showed significant upregulation of glutamate decarboxylase (gadB), GABA transaminase (gabT), and succinic semialdehyde dehydrogenases (gabD1 and GabD2) (Supplementary Figure S1A–D). Notably, the succinate-producing enzyme gabD showed an interesting pattern, where gabD1 and gabD2 were differentially induced under acidic and oxidative stress, suggesting context-specific roles for these paralogs.

We focused on *Mtb*GadB, the first and rate-limiting enzyme of the GABA shunt pathway, which was upregulated under both acidic and oxidative stresses. Consistent with transcript levels, *Mtb*GadB protein levels increased nearly 2-fold under both stresses (Fig. 1A, 1B). To determine the functional impact, we measured GABA production in stressed *Mtb* cultures. Intrabacterial GABA increased by ∼2-fold under acidic and ∼3-fold under oxidative stress, as shown by a modified Berthelot assay (Fig. 1C). Given reports of GABA release by some gut microbiota. (31), we also quantified extracellular GABA. LC-MS analysis demonstrated enhanced GABA in the culture supernatant of *Mtb* exposed to acidic conditions compared to control, further quantified by the modified Berthelot assay (32) (Fig. 1D). These results confirm the activation of GABA shunt pathway, with *Mtb*GadB-driven GABA accumulation under stress, supporting a role for this pathway in *Mtb* metabolic adaptation.

**Figure 1:**
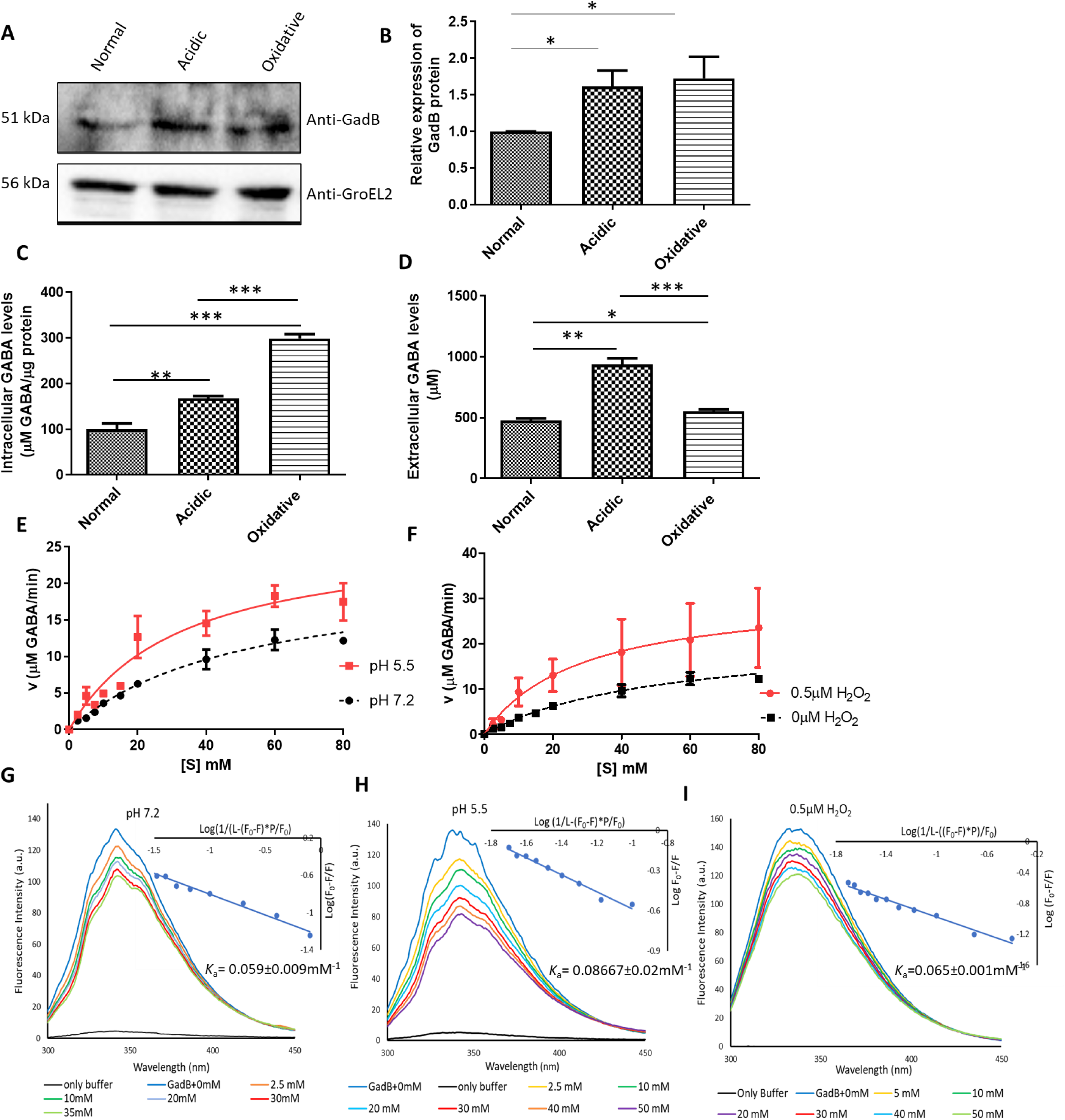
Expression and catalytic efficiency of *Mtb*GadB are increased under acidic and oxidative stress in *Mycobacterium tuberculosis*. (A) Immunoblots of *Mtb*GadB protein under acidic and oxidative microbicidal stress conditions in *Mtb* H37Rv cell lysates. Equal amounts of lysate (Bradford-normalized) were loaded, with GroEL2 serving as the loading control. (B) Immunoblot-derived quantification of *Mtb*GadB, normalized to GroEL2 and presented relative to the normal condition. (C) Bar graph showing quantification of the total amount of intracellular GABA produced under stress, as quantified by the modified Berthelot assay and normalized to total protein concentration. (D) Bar graph showing extracellular GABA levels released under acidic and oxidative stress. The standard GABA curve was used as the reference for quantification in panels C and D. (E) Michaelis-Menten plots of *Mtb*GadB at pH 5.5 and pH 7.2 (control); (F) Michaelis-Menten plots of *Mtb*GadB in the presence (0.5 μM) and absence of H_2_O_2_ (control) (G-I) Representative fluorescence spectra of quenching of emission of *Mtb*GadB with L-glutamate under different conditions. Modified Stern-Volmer equations with the *K*_a_ values of L-glutamate at (G) pH 7.2, (H) pH 5.5, and (I) 0.5 µM of H_2_O_2_. All the experiments were performed in biological triplicate. Statistical significance was determined using an unpaired Student’s t-test. The p-values are denoted as ‘***’ p ≤0.0005; ‘**’ p ≤ 0.005, ‘*’ p≤ 0.05, while ns denote non-significant values.

To further investigate how *Mtb*GadB behaves enzymatically under these two independent yet related stresses, we cloned, expressed, and purified recombinant *Mtb*GadB (Supplementary Fig. S2A–C) and compared its kinetic parameters under acidic and oxidative conditions. To test if the recombinant *Mtb*GadB was active, reactions with L-glutamate as a substrate were performed, and the production of GABA was confirmed using HPLC (Supplementary Figure S2D–F). Under identical conditions, the end-point production of GABA was measured using a modified Berthelot assay (detailed in methodology) to determine kinetic parameters (*K*_m_ and *V*_max_) and catalytic efficiencies (*k*_cat_/*K*_m_). Under acidic conditions, a marked decrease in *K*_m_ for L-glutamate, with a moderate increase in turnover (*k*_cat_), led to ∼2-fold higher catalytic efficiency relative to neutral pH. Under oxidative conditions, both an increase in turnover and a reduction in *K*_m_ contributed to a ∼2.5-fold enhancement in catalytic efficiency (Fig. 1E, 1F; Table 1).

**Table 1:** Kinetic parameters of *Mtb*GadB at pH 7.2, pH 5.5 and 0.5µM of H_2_O_2_

| Kinetic parameter | pH 7.2 | pH 5.5 | 0.5μM H<sub>2</sub>O<sub>2</sub> |
| --- | --- | --- | --- |
| $V_{\max}$ (μM GABA/min) | 21.84±3.576 | 26.75±3.77 | 31.79±9.747 |
| $K_m$ (mM) | 51.02±15.54 | 32.36±10.20 | 28.76±19.67 |
| $k_{\text{cat}}$ (sec<sup>-1</sup>) | 0.145±0.023 | 0.178±0.025 | 0.211±0.06 |
| Catalytic efficiency $k_{\text{cat}}/K_m$ (sec<sup>-1</sup> mM<sup>-1</sup>) | 0.0028±0.0014 | 0.0055±0.0024 | 0.007±0.003 |
| $K_a$ (mM<sup>-1</sup>) | 0.059±0.009 | 0.087±0.02 | 0.065±0.001 |

Because a lower *K*_m_ indicates higher substrate affinity, we also determined the association (binding) constants (*K*_a_) for L-glutamate binding to *Mtb*GadB under acidic and oxidative conditions using fluorescence spectroscopy. The Stern-Volmer equation was applied to calculate the *K*_a_ values (Table 1). *K*_a_ was higher under both acidic and oxidative conditions than in the control, supporting the observed lower *K*_m_ values and indicating increased substrate affinity (Fig. 1G–I).

Collectively, these data indicate that acidic and oxidative stresses enhance the catalytic efficiency of *Mtb*GadB by increasing substrate binding, thereby increasing GABA production.

### C-terminal domain orchestrates catalytic efficiency and pH-dependent activity of *Mtb*GadB

Our *in vitro* enzymatic assays demonstrated that *Mtb*GadB exhibits increased catalytic efficiency under both acidic and oxidative conditions. We therefore sought to identify the structural features of *Mtb*GadB that underlie this enhancement, as such features are likely to contribute to intracellular persistence during infection.

To address this, we generated a three-dimensional model of *Mtb*GadB using the *E.coli* Gad crystal structure as a template, since the crystal structure of *Mtb*GadB is not yet available. Computational analyses have demonstrated pH-dependent conformational changes in *E.coli* Gad (33). In addition, mutational studies of *Lactobacillus* Gad, based on homology modeling using *E.coli* Gad, highlighted the critical role of the C-terminal region in pH-dependent catalysis (34). Together, these findings motivated us to use *E.coli* Gad as a reliable template for modeling *Mtb*GadB. The predicted structure of *Mtb*GadB displayed the canonical PLP-dependent decarboxylase fold, comprising an N-terminal region (residues 1–58), a catalytic domain (58–346), and a C-terminal region (348–460). The C-terminal region of *Mtb*GadB differs markedly from that of *E.coli* Gad. In *E.coli*, it forms a stable α-helix-loop, whereas in *Mtb* it adopts a flexible loop (Figure 2A). Despite this structural difference, analysis of the *Mtb*GadB model suggests that the C-terminal region contributes to protein stability, with residues Asp445, Lys446, Phe453, and His460 playing key roles. Given that *Mtb* encounters acidic environments during macrophage infection, we hypothesized that the C-terminal region may similarly regulate stress-responsive activity, and that its removal could affect the pH-dependent stability of the protein. To test this, we generated a C-terminal truncation mutant lacking residues Val447–His460 (*Mtb*GadBΔCt). To distinguish regulatory effects from catalytic defects and to confirm that regulation arises from the C-terminal domain (Val447-His460) (CTD) rather than the catalytic domain, we generated a control catalytic mutant using site-directed mutagenesis. In this mutant, the conserved active-site residues His276 and Lys277 (Figure 2B) were substituted with alanine (*Mtb*GadBSDM), while the CTD was left intact. Both mutants were purified under native conditions for biochemical characterization (Supplementary Figure S3).

**Figure 2:**
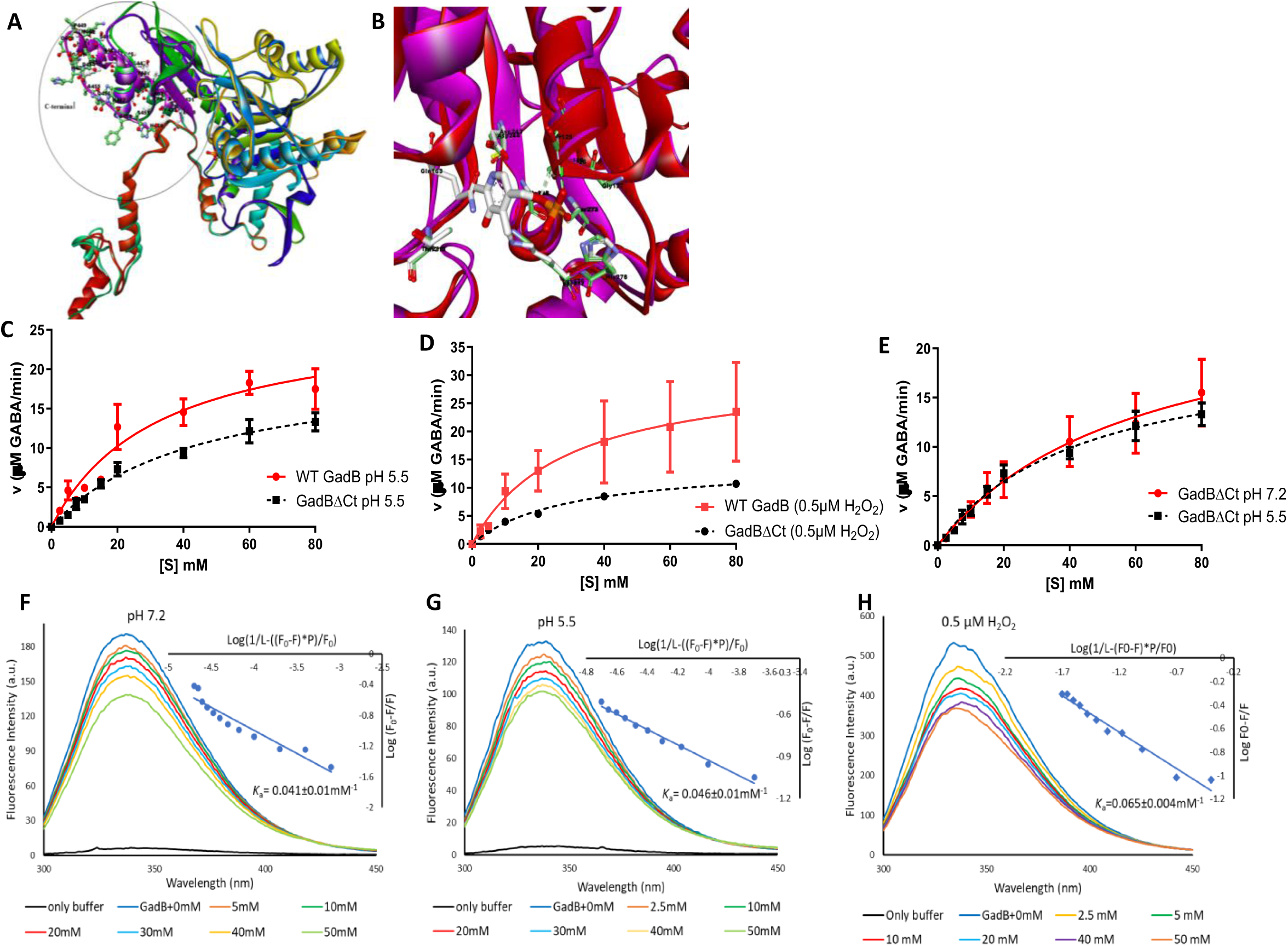
Deletion of the C-terminal domain reduced the catalytic efficiency of *Mtb*GadB under acidic and oxidative stresses. (A) Cartoon representation of the superimposition of C - terminal domains of *Mtb*GadB and *E.coli* GadB. (B) Cartoon representation of the catalytic site of *Mtb*GadB highlighting a highly conserved lysine residue (Lys277), which forms a Schiff base with the PLP cofactor, and other residues including Asp244, Gly127, Ser128, Ser129, Ser274, and His276. (C) Michaelis Menten plots of *Mtb*GadB and *Mtb*GadBΔCt at pH 5.5; (D) Michaelis Menten plots of *Mtb*GadB and *Mtb*GadBΔCt in the presence of 0.5 μM H_2_O_2_ (E) Michaelis Menten plot of *Mtb*GadBΔCt at pH 5.5 and pH 7.2 (F-H) Representative fluorescence spectra of quenching of emission of *Mtb*GadBΔCt with L-glutamate under different conditions. Modified Stern-Volmer plots for the fluorescence quenching of the *Mtb*GadBΔCt at (F) pH 7.2, (G) pH 5.5, and (H) 0.5µM of H_2_O_2_. All the experiments were performed in biological triplicates.

*In vitro* enzymatic assays were performed with purified *Mtb*GadBΔCt and *Mtb*GadBSDM, and kinetic parameters were evaluated as described previously. *Mtb*GadBΔCt showed reduced catalytic turnover and increased *K*_m_ under acidic conditions, resulting in ∼2-fold decrease in catalytic efficiency compared to wild-type *Mtb*GadB (Figure 2C). This increase in *K*_m_ was further confirmed by fluorescence spectroscopy, which demonstrated that the binding constant (*K*_a_) of *Mtb*GadBΔCt to L-glutamate was reduced to half that of wild-type *Mtb*GadB under acidic conditions (Figure 2G). To investigate the role of the CTD in pH-dependent catalytic efficiency, we compared the catalytic parameters of *Mtb*GadBΔCt at acidic (pH 5.5) and neutral (pH 7.2) conditions. *Mtb*GadBΔCt displayed similar *k*_cat_/*K*_m_ and *K*_a_ values at both pH levels, indicating that deletion of the CTD not only decreased catalytic efficiency but also abolished pH-dependent activity, likely by impairing L-glutamate binding (Figure 2E-G, Table 2). To further confirm that the pH-dependent increase in catalytic efficiency is mediated by the CTD, rather than the catalytic domain, we performed kinetic assays with the catalytic mutant *Mtb*GadBSDM under acidic conditions. *Mtb*GadBSDM exhibited a ∼2-fold decrease in both catalytic efficiency and binding constant (*K*_a_) compared to wild-type *Mtb*GadB (Supplementary Figure S4A, S4F-G). However, *Mtb*GadBSDM retained higher catalytic efficiency at acidic pH than at neutral pH, mirroring wild-type *Mtb*GadB (Supplementary Figure S4C). This preservation of pH-dependent behavior in *Mtb*GadBSDM confirms that the catalytic domain is not responsible for pH sensing and underscores the role of the CTD in mediating pH-dependent increase in catalytic efficiency of *Mtb*GadB. Tables S3 and S4 list the kinetic parameters of *Mtb*GadBΔCt and *Mtb*GadBSDM.

**Table 2:** Catalytic efficiencies (*k*_cat_/*K*_m_) and association constants (*K*_a_) of *Mtb*GadBΔCt and *Mtb*GadBSDM under acidic and oxidative conditions.

| Enzyme | Kinetic parameter | pH 7.2 | pH 5.5 | 0.5μM H<sub>2</sub>O<sub>2</sub> |
| --- | --- | --- | --- | --- |
| *Mtb*GadBΔCt | $k_{\text{cat}}/K_m$ (sec<sup>-1</sup> mM<sup>-1</sup>) | 0.0028±0.001 | 0.003±0.001 | 0.003±0.001 |
| | $K_a$ (mM<sup>-1</sup>) | 0.041±0.01 | 0.046±0.01 | 0.065±0.004 |
| *Mtb*GadBSDM | $k_{\text{cat}}/K_m$ (sec<sup>-1</sup> mM<sup>-1</sup>) | 0.0021±0.001 | 0.003±0.001 | 0.0031±0.0016 |
| | $K_a$ (mM<sup>-1</sup>) | 0.053±0.004 | 0.062±0.007 | 0.056±0.01 |
| WT *Mtb*GadB | $k_{\text{cat}}/K_m$ (sec<sup>-1</sup> mM<sup>-1</sup>) | 0.0028±0.0014 | 0.0055±0.0024 | 0.007±0.003 |
| | $K_a$ (mM<sup>-1</sup>) | 0.059±0.009 | 0.08667±0.02 | 0.065±0.001 |

We next focused on the possible impact of the CTD of *Mtb*GadB on catalysis under oxidative conditions. *In vitro* enzymatic assays were performed with *Mtb*GadBΔCt and *Mtb*GadBSDM in the presence of 0.5 μM H_2_O_2_, as previously described. *Mtb*GadBΔCt exhibited reduced catalytic turnover (*k*_cat_) but maintained a *K*_m_ similar to wild-type *Mtb*GadB, resulting in ∼2-fold decrease in catalytic efficiency (Figure 2D). This similarity in *K*_m_ between *Mtb*GadBΔCt and wild-type *Mtb*GadB under oxidative conditions was further supported by binding constant (*K*_a_) measurements using fluorescence spectroscopy (Figure 2H, Table 2). These results indicate that deletion of the CTD impairs catalytic turnover without affecting substrate binding affinity during oxidative stress. Since we observed the role of CTD in pH-dependent regulation of catalytic efficiency, we further examined its involvement under oxidative conditions. Kinetic assays using *Mtb*GadBSDM, which retains a functional CTD, revealed a 50% decrease in catalytic efficiency compared to wild-type *Mtb*GadB, suggesting that His276 and Lys277 are important for activity under oxidative stress (Supplementary Figure S4B, S4H).

Together, these results show that the C-terminal domain of *Mtb*GadB functions as a regulatory element that confers pH-dependent increase in catalytic efficiency, primarily by enhancing substrate binding at acidic pH. In contrast, the catalytic domain governs enzymatic turnover but is not involved in sensing pH changes. Under oxidative conditions, the C-terminal domain supports catalytic efficiency rather than regulating substrate affinity, suggesting condition-specific functional contributions.

### C-terminal domain of *Mtb*GadB regulates intrabacterial GABA production and ROS reduction shaping the bacterial stress-adaptive metabolic state

Our comparative enzyme kinetic analyses showed that deletion of the C-terminal domain (CTD) markedly reduces the catalytic efficiency of *Mtb*GadB under acidic and oxidative conditions. Importantly, these findings indicate that the CTD plays a critical role in regulating the enzyme’s pH-dependent activity. To assess the physiological relevance of this observation, we expressed wild-type and mutant *Mtb*GadB constructs in the pathogenic *Mtb H37Rv* (Supplementary Figure S5). Strains overexpressing wild-type *Mtb*GadB, a catalytic mutant (*Mtb*GadBSDM), or a C-terminal truncation mutant (*Mtb*GadBΔCt) were subjected to acidic and oxidative stresses. Intrabacterial GABA levels were quantified by NMR and expressed as the ratio of GABA produced under stress versus control conditions. Under acidic conditions, the ratio of GABA levels at pH 5.5 relative to pH 7.2 was higher in the wild-type overexpression strain (*Mtb::Mtb*GadB) compared to the vector control, indicating enhanced acid-responsive GABA production. As expected, the catalytic mutant (*Mtb::Mtb*GadBSDM) produced lower GABA levels than the wild-type. However, it retained an increased GABA ratio at low pH, suggesting that pH-dependent regulation is largely preserved despite reduced catalytic activity. In contrast, the CTD truncation mutant (*Mtb::Mtb*GadBΔCt) not only exhibited reduced GABA production but also showed a GABA ratio close to unity, demonstrating a loss of pH responsiveness. These results establish that the CTD is essential for pH-dependent GABA production (Figure 3A). A similar trend was observed under oxidative stress as well. *Mtb::Mtb*GadB increased GABA levels compared to the vector control, whereas both *Mtb::Mtb*GadBSDM and *Mtb::Mtb*GadBΔCt produced lower amounts of GABA (Figure 3B).

**Figure 3:**
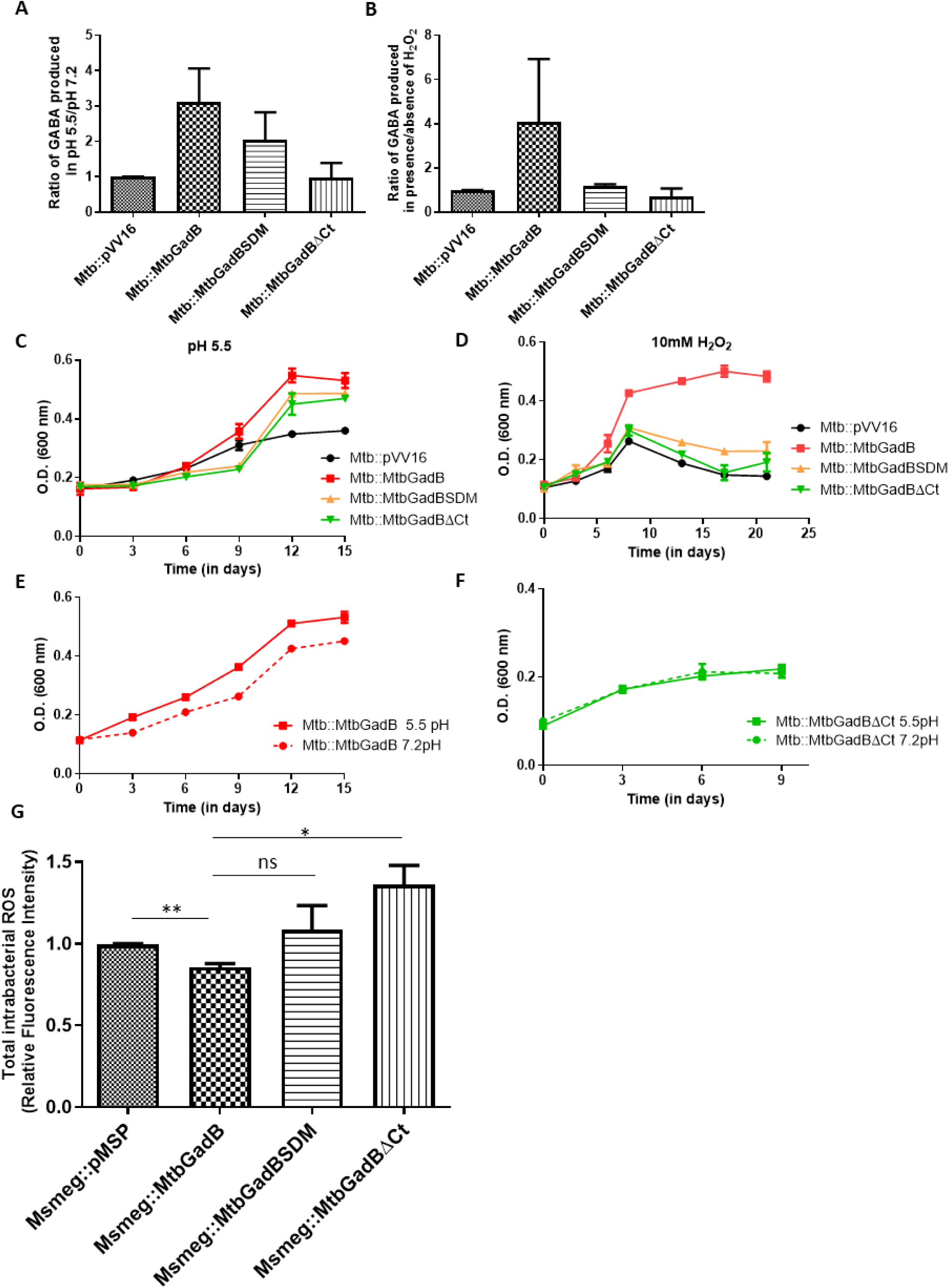
Deletion of C-terminal domain impaired GABA production and stress tolerance in *Mtb*. (A) Ratio of GABA produced at pH 5.5 to pH 7.2 (B) Ratio of GABA produced in the presence and absence of H_2_O_2_. The data were first normalized to the total protein concentration, followed by normalization to the vector control (*Mtb*::pVV16), and reported as ratios. (C) Time-growth plots of *Mtb::Mtb*GadB, *Mtb::Mtb*GadBSDM and *Mtb::Mtb*GadBΔCt in acidic pH (D) Time-growth plots of *Mtb::Mtb*GadB, *Mtb::Mtb*GadBSDM, and *Mtb::Mtb*GadBΔCt in H_2_O_2_. *Mtb*::pVV16 was used as a control. (E) Time-growth plots of *Mtb::Mtb*GadB in acidic and neutral pH. (F) Time-growth plot of *Mtb::Mtb*GadBΔCt in acidic and neutral pH. (G) Quantification of intra-bacterial ROS by Fluorescence spectroscopy. Unstained bacteria were used as a control, and the data were normalized to the *Msmeg*::pMSP vector control. All the experiments were performed in biological triplicate. Statistical significance was determined using an unpaired Student’s t-test. The p-values are denoted as ‘***’ p ≤0.0005; ‘**’ p ≤ 0.005, ‘*’ p≤ 0.05, while non-significant values are denoted by ns.

The functional consequences of these biochemical differences were evident in *in vitro* growth assays. *Mtb::Mtb*GadB displayed enhanced growth under acidic stress relative to the control strain *Mtb*::pVV16 (Figure 3C). *Mtb::Mtb*GadB exhibited improved growth at acidic pH compared to neutral pH, whereas the CTD truncation mutant (*Mtb::Mtb*GadBΔCt) displayed similar growth profiles under both conditions, indicating a loss of pH- enhanced growth advantage (Figures 3E–F). Under oxidative stress, *Mtb::Mtb*GadB showed improved growth compared to *Mtb::Mtb*GadBΔCt as well as *Mtb::Mtb*GadBSDM (Figure 3D). As we understand that GABA production by *Mtb*GadB consumes a proton (H⁺), thereby buffering intracellular pH and reducing conditions that favor ROS amplification, indirectly contributing to ROS quenching (35), we measured intrabacterial ROS using CM-H_2_DCFDA dye in *Mycolicibacterium smegmatis* (*Msmeg*) strains overexpressing wild-type *Mtb*GadB, *Mtb*GadBSDM and *Mtb*GadBΔCt, under oxidative stress. Consistent with their *Mtb*GadB enzymatic efficiencies, intracellular ROS in *Msmeg::Mtb*GadB was 20% less than *Msmeg*::pMSP (control). In contrast, *Msmeg::Mtb*GadBΔCt had elevated ROS accumulation relative to *Msmeg::Mtb*GadB (Figure 3G). Collectively, these findings suggest that *Mtb*GadB enhances the survival of *Mtb* under acidic and oxidative stress by promoting GABA production and limiting ROS accumulation. Crucially, the C-terminal domain is indispensable for this stress-adaptive function, as its deletion compromises both pH-dependent regulation and the ability of *Mtb*GadB to mitigate intracellular oxidative stress.

### CTD-regulated GABA-producing metabolic state of *Mtb* blocks endosomal maturation and dampens oxidative stress during infection

After establishing that *Mtb::Mtb*GadB aids bacterial growth under acidic and oxidative stresses, we next investigated the intracellular fate of these biochemically distinct strains during infection. We hypothesized that the strain overexpressing wild-type (wt) *Mtb*GadB, due to its enhanced conversion of glutamate to GABA, would facilitate proton quenching, thereby contributing to neutralization of the acidic environment of phagolysosomes. Furthermore, we proposed that increased *Mtb*GadB and GABA production would serve as an indirect defense mechanism against oxidative stress, reducing macrophage ROS generation during infection. In contrast, strains overexpressing *Mtb*GadBSDM and *Mtb*GadBΔCt would lack these protective effects. To test our hypothesis, we used mCherry-tagged *Msmeg*::pMSP, *Msmeg*::*Mtb*GadB, *Msmeg*::*Mtb*GadBSDM, and *Msmeg*::*Mtb*GadBΔCt for live cell imaging experiments.

THP-1 macrophages were infected with the indicated *Msmeg* strains (MOI 1:20) and stained at 6 hours post-infection (hpi) for quantification of LysoTracker intensity by confocal microscopy. Line intensity profile analysis demonstrated overlapping spatial distribution of *Msmeg* and LysoTracker signals across all strains, consistent with similar Manders’ M1 coefficients (Figure 4A). However, LysoTracker intensity at bacteria-associated regions was reduced in the *Msmeg*::*Mtb*GadB strain compared to the vector control and mutant strains (Figure 4B-D, Supplementary Figure S6). These results indicate that although the strains do not differ in their ability to localize to acidified compartments (similar M1 coefficient), they do exhibit differential acidification of these compartments, as reflected in their LysoTracker intensity profiles. Notably, infection with the strain expressing the most catalytically efficient wild-type *Mtb*GadB (*Msmeg*::*Mtb*GadB) resulted in the greatest neutralization of acidic compartments.

**Figure 4:**
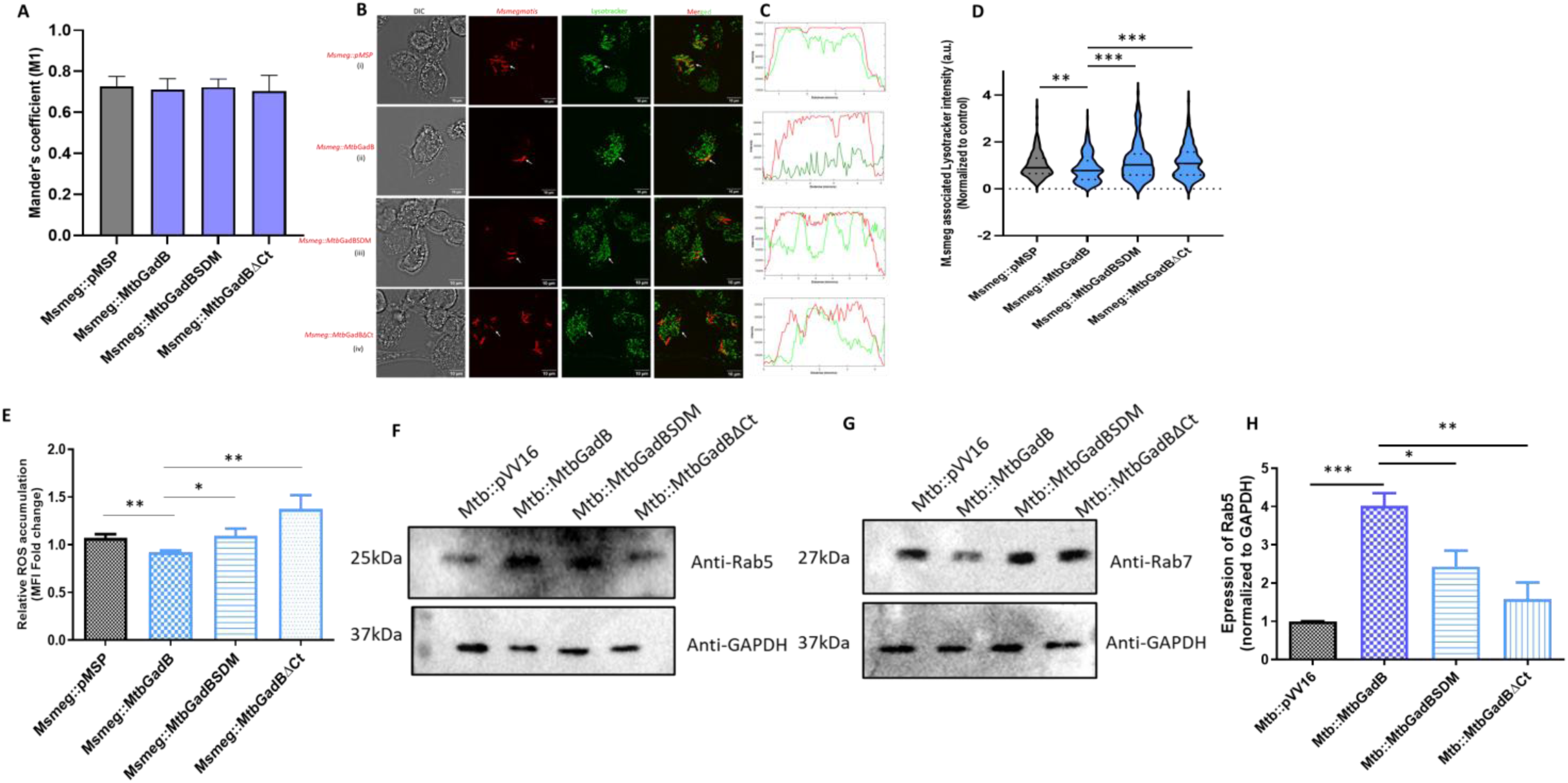
CTD-regulated GABA-producing metabolic state of *Mtb* blocks endosomal maturation and dampens oxidative stress during infection: (A) Bar-graph representing Mander’s colocalization coefficient (M1) of different strains of *Msmegmatis* with Lysotracker stained compartments (B) Confocal microscopy of Lysotracker-stained THP-1 macrophages infected with different cherry-tagged *Msmeg* strains showing localization with acidified compartments. Acidified compartments are seen as green fluorescence, and *Msmeg* is seen in red fluorescence. Bacteria localised with acidified compartments are seen in yellow fluorescence. (C) The plot profiles of representative single-cell images of insets (i-iv) for lysotracker positive compartments (green line) and *Msmeg* (red line) were generated using ImageJ software. (D) Violin plots showing the mean Lysotracker intensity associated with individual bacterial region of interests (ROIs). Individual data points represent the background-corrected lysotracker intensity associated with a bacterial ROI of the particular strain. Data was normalized to control (*Msmeg*::pMSP) and represented as *Msmeg* asssociated Lysotracker intensity. (E) Bar graph showing quantification of Median Fluorescence Intensity (MFI) of CM-H_2_DCFDA dye representing intracellular ROS measured by flow cytometry. Data were normalized to the PMA-treated group and reported as fold change. (F-G) Immunoblots showing protein levels of Rab5 (G) and Rab7 (H) under infection conditions in THP-1 cell lysates. Equal amounts (as measured by Bradford assay) of lysate were loaded in each well, and GAPDH was used as an internal control. (H) Bar graph showing the ratio of Rab5/Rab7 expression by immunoblot-based quantification of expression, normalized to GAPDH, and represented as relative ratios compared to vector control. All the experiments were performed at least in biological triplicate. Statistical significance was determined using an unpaired Student’s t-test. The p-values are denoted as ‘***’ p ≤0.0005; ‘**’ p ≤ 0.005, ‘*’ p≤ 0.05., while non-significant values are denoted by ns.

Similarly, intracellular ROS levels were quantified 6 hours post-infection using CM-H_2_DCFDA staining and flow cytometry. Macrophages infected with *Msmeg*::*Mtb*GadB exhibited a ∼20% reduction in ROS levels compared to the vector control (Figure 4E, Supplementary Figure S7). Consistent with their reduced capacity to produce GABA, infections with *Msmeg*::*Mtb*GadBSDM and *Msmeg*::*Mtb*GadBΔCt resulted in higher ROS accumulation relative to *Msmeg*::*Mtb*GadB.

Because endosomal acidification and ROS levels are indicators of endosomal maturation (36), we assessed the expression of early and late endosome markers Rab5 and Rab7, respectively, during infection with these strains. The Rab5/Rab7 ratio served as a metric for endosomal maturation, reflecting the shift from early (Rab5-positive) to late (Rab7-positive) endosomes. We observed that the strain expressing wild-type *Mtb*GadB (*Mtb::Mtb*GadB) showed the highest Rab5/Rab7 ratio, followed by *Mtb*::*Mtb*GadBSDM. In contrast, the *Mtb*::*Mtb*GadBΔCt mutant displayed Rab5/Rab7 ratios similar to control infections, indicating the least impairment in endosomal maturation. Infection with *Mtb*::*Mtb*GadB led to a ∼4-fold increase in the Rab5/Rab7 ratio compared to the vector control (*Mtb*::pVV16), indicating substantial inhibition of endosomal maturation by this strain (Figure 4F-H). These findings demonstrate that *Mtb*GadB expression decreases host endosomal maturation, whereas loss of CTD and the accompanying reduction in catalytic efficiency diminish this effect.

Overall, our results suggest that *Mtb*GadB-dependent GABA production dampens host-induced intracellular acidic and oxidative stresses and delays endosomal maturation during infection.

### The CTD of *Mtb*GadB is required for M2 macrophage polarisation and intracellular persistence during infection

Given that infection with *Msmeg:*:*Mtb*GadB clearly decreases ROS levels and phagosomal maturation, a classical feature of M2-macrophages, and considering the established immunometabolic role of GABA in various intracellular infections (37–39), it was essential to examine the total intracellular GABA pool, following infection with these biochemically distinct strains and the polarisation state of the infected macrophages.

The intracellular GABA pool following infection was quantified using the modified Berthelot assay. THP-1 macrophages infected with *Mtb::Mtb*GadB exhibited significantly higher intracellular GABA levels compared to control infections (*Mtb*::pVV16). In contrast, infections with *Mtb::Mtb*GadBSDM and *Mtb::Mtb*GadBΔCt resulted in markedly reduced intracellular GABA levels compared to *Mtb::Mtb*GadB (Figure 5A), indicating that the ability of *Mtb* to synthesize GABA affects the total GABA pool of macrophages during infection.

**Figure 5:**
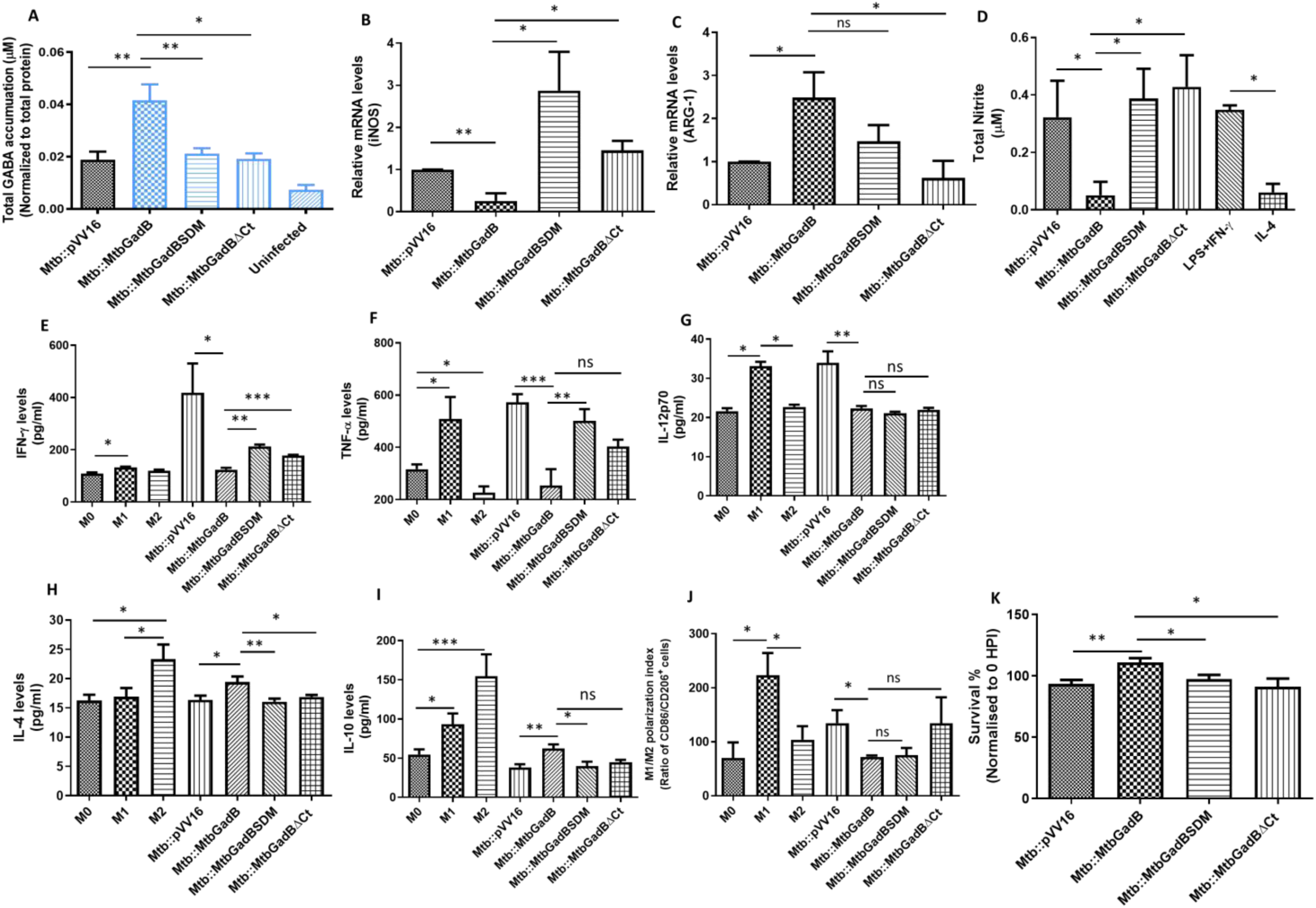
The C-terminal domain of *Mtb*GadB is required for M2 macrophage polarisation and intracellular persistence during infection. (A) Bar graph showing the total amount of GABA accumulated within THP-1 macrophages upon infection with different strains of *Mtb*. Uninfected THP-1 macrophages served as controls. Data was normalized to the total protein concentration of each sample. (B-D) Transcript levels of (A) iNOS and (B) Arginase-1 (Arg-1) in THP-1 macrophages were quantified by qRT-PCR at 24 hpi. *Mtb*::pVV16 infected macrophages were used as control. GAPDH was used as an internal control. (D) Bar graph showing total nitrite levels in the culture supernatant of infected THP-1 macrophages, as measured by the Griess assay. LPS+IFN-y and IL-4 treated macrophages were used as control for M1 and M2 polarized states respectively. A standard plot of nitrite was used for the quantification of total nitrite. (E-I) ELISA based quantification of different cytokines (pro-inflammatory (E) [IFN-γ], (F) [TNF-α]), (G) [IL-12p70] and anti-inflammatory (H) [IL-4], (I) [IL-10]) in the culture supernatant of THP-1 macrophages at 24 hours post-infection (hpi). Uninfected and M1/M2 polarized macrophages were used as controls. (J) Flow-cytometry based analysis of CD86 and CD206 positive population of macrophages upon infection with different strains of *Mtb*. Data were represented as a ratio of CD86/CD206-positive population. M1/M2 polarized macrophages were used as controls. (K) Bar-graph showing % survival of *Mtb* in THP-1 macrophages. Infected macrophages were lysed and plated on 7H10 agar plates, and CFU was calculated at 24 hpi. Data was normalized to a CFU count at 0 hpi and represented as % survival. Error bars represent standard deviation. All the experiments were performed at least in biological triplicate. Statistical significance was determined using an unpaired Student’s t-test. The p-values are denoted as ‘***’ p ≤0.0005; ‘**’ p ≤ 0.005, ‘*’ p≤ 0.05., while non-significant values are denoted by ns.

We next examined different phenotypic markers of macrophage polarisation in THP-1 macrophages infected with *Mtb::Mtb*GadB, *Mtb::Mtb*GadBSDM, and *Mtb::Mtb*GadBΔCt. Infection with *Mtb::Mtb*GadB led to an approximately 50% reduction in iNOS expression and nitrite production, as well as a ∼2-fold increase in arginase-1 (Arg-1) expression. These changes indicate decreased generation of reactive nitrogen species (RNS) and a shift away from classical M1 macrophage activation (Figure 5B-D). Consistent with this, *Mtb::Mtb*GadB infection reduced the levels of pro-inflammatory cytokines (IFN-γ, TNF-α, and IL-12p70) and increased anti-inflammatory cytokines (IL-4 and IL-10) (Figure 5E-I). Flow cytometry further revealed reduced CD86 expression and a lower CD86/CD206 ratio, indicating polarisation toward an M2-like phenotype (Figure 5J, Supplementary Figure S8). As expected, these effects were less pronounced in macrophages infected with *Mtb::Mtb*GadBSDM or *Mtb::Mtb*GadBΔCt, suggesting that the GABA-producing capacity of *Mtb* drives this immunomodulation and creates a niche favourable for intracellular mycobacterial survival within THP-1 macrophages. Therefore, we assessed colony-forming unit (CFU) enumeration at specified time points post-infection. Results showed that *Mtb::Mtb*GadB had the highest CFU count, with ∼1.5-fold increase in intracellular survival relative to the vector control. In contrast, *Mtb::Mtb*GadBSDM showed moderate, and *Mtb::Mtb*GadBΔCt the lowest, CFU counts, indicating enhanced bacterial clearance by macrophages (Figure 5K).

Collectively, these results show that the CTD of *Mtb*GadB orchestrates the accumulation of GABA in macrophages during infection, polarising them towards an M2-phenotype, dampening the *Mtb* clearance responses.

## Discussion

Intracellular survival of *Mtb* relies on integrating responses to diverse host-imposed stresses and reshaping the immune environment (10, 40). Traditionally, acid resistance and redox homeostasis were viewed as parallel processes, but our results indicate metabolic coordination between these stresses and a functional linkage to host immunomodulation via glutamate decarboxylase (*Mtb*GadB). Our study shows that *Mtb*GadB is not merely stress-responsive in *Mtb*; it is optimized for the intracellular environment, especially through condition-specific enhanced substrate binding. Whereas non-pathogenic mycobacteria such as *Mycolicibacterium smegmatis* (*Msmegmatis*) employ this enzyme against environmental stresses, pathogenic mycobacteria like *Mtb* may have evolved to exploit it to counteract host-induced bactericidal pressures. Glutamate decarboxylation consumes intracellular protons, aiding pH homeostasis, while the GABA shunt influences redox balance through NAD⁺/NADH regulatory pathways (24, 35). Further, enhanced GABA may mitigate oxidative stress by lowering intracellular ROS and stabilizing redox homeostasis (9). Notably, our observation that both acidic and oxidative stress upregulate *Mtb*GadB and the GABA shunt pathway (Figure 1) supports a functional coupling between proton consumption, GABA-mediated antioxidant effects, and redox balance, positioning *Mtb*GadB at the nexus of multiple stress adaptation pathways. Furthermore, differential expression of gabD1 and gabD2 under acidic and oxidative stress (Supplementary Figure S1) implies that *Mtb* may channel GABA-derived carbon flux through alternative metabolic branches depending on the stress encountered. This regulation fine-tunes redox homeostasis and energy metabolism, revealing metabolic flexibility within the GABA shunt pathway.

Structural and mutational analyses identify the C-terminal domain (CTD) as a key determinant of *Mtb*GadB function under stress. Deleting the CTD reduced catalytic efficiency (*k*_cat_/*K*_m_) and abolished pH-dependent variation in activity, indicating the CTD is essential for optimal, pH-responsive enzyme performance (Figure 2). This likely occurs through effects on conformational dynamics or substrate access, rather than acting as a direct catalytic determinant. Interestingly, the loss of the CTD also led to reduced enzymatic activity under oxidative conditions, suggesting that this domain may play a broader role in maintaining stress-responsive enzyme function. Since acidic and oxidative stress can cause structural changes and alter residue protonation (41), the CTD may stabilize the active conformation of *Mtb*GadB, linking pH sensing with oxidative stress adaptation. This is relevant for pyridoxal phosphate (PLP)-dependent enzymes, whose activity depends on the active-site microenvironment and is influenced by redox-driven conformational changes (42, 43), further supporting the CTD’s role in maintaining catalytic competence under variable conditions. Conversely, mutation of active-site residues mainly reduced turnover while preserving stress responsiveness, supporting a model in which the CTD enables dynamic conformational changes to optimize function under stress. Comparative analysis of *Mtb*GadB from that of *Msmegmatis*, a non-pathogenic species, reveals greater sequence variability and structural flexibility in its C-terminal region. This likely reflects different evolutionary pressures, where environmental mycobacteria need adaptability, while pathogenic *Mtb* requires stringent regulation to survive in a defined but hostile host niche. Overall, the *Mtb*GadB C-terminal region appears to have specialized in pathogenic mycobacteria, evolving from a flexible structure into a regulatory domain that integrates pH sensing and oxidative stress adaptation, a mechanism that may underlie *Mtb*’s persistence and resilience within the host.

Deletion of CTD resulted in decreased GABA production and mitigated adaptation to acidic and oxidative stresses (Figure 3). Further, infection with *Msmeg::Mtb*GadB decreased phagosomal maturation and reduced ROS compared to *Msmeg::*pMSP as well as *Msmeg*::*Mtb*GadBΔCt (Figure 4), indicating that GadB-mediated metabolic state of mycobacteria influences the stress-adaptive response against acidic and oxidative stresses within THP-1 macrophages. Further, beyond its established role in bacterial physiology, our data demonstrate that stress-induced *Mtb*GadB-mediated GABA production significantly influences host–pathogen interactions. We detected GABA in the culture supernatant of *Mtb* exposed to *in vitro* acidic and oxidative stress, despite no reported antiporters in *Mtb* (Figure 1). In high-GABA-producing strains, GABA may leak or be exported via unknown mechanisms. Using *Mtb* strains overexpressing *Mtb*GadB (*Mtb::Mtb*GadB), we observed that infected macrophages showed markedly higher intracellular GABA, whereas infections with *Mtb::Mtb*GadBSDM and *Mtb::Mtb*GadBΔCt did not (Figure 5A). This suggests that the ability of *Mtb* to synthesize GABA affects the total GABA pool during infection. Whether this increase results from bacterial GABA leakage, or GABA release due to bacterial lysis, or infection-induced host GABA synthesis remains unclear. Dissecting the relative contributions of bacterial- versus host-derived GABA will require targeted approaches, such as host GAD inhibition or isotope tracing, and represents a key area for future research.

Our study demonstrates that *Mtb* strains overexpressing *Mtb*GadB and producing increased GABA suppress multiple pro-inflammatory responses, driving macrophage polarisation towards an M2-like phenotype (Figure 5). Although we observed increased expression of GABA transporters (GATs) and GABA receptor subunit (GABRA2) in THP-1 infected with *Mtb::Mtb*GadB as compared to *Mtb::Mtb*GadBSDM and *Mtb::Mtb*GadBΔCt (Supplementary figure S9), the specific receptor-mediated mechanisms in this context remain to be elucidated. These immunomodulatory effects require an intact CTD, directly connecting stress-responsive enzyme activity with host immunomodulation. Notably, the diminished survival of the C-terminal truncation mutant highlights the essential role of *Mtb*GadB-mediated metabolism in infection, linking stress adaptation and immunomodulation to bacterial persistence in macrophages.

In summary, this study identifies the C-terminal domain of *Mtb*GadB as a pivotal structural and functional determinant for enhanced catalytic efficiency under acidic and oxidative conditions, as well as governing pH-dependent enzymatic activity. This critical role directly contributes to intracellular stress adaptation and immunomodulatory processes, thereby enhancing the survival capacity of *Mtb* within host cells during infection. Figure 6 represents a proposed model of the impact of stress-adaptive metabolic state of *Mtb*::*Mtb*GadB and *Mtb*::*Mtb*GadBΔCt on the intracellular survival of *Mtb*. Notably, the marked structural divergence between the C-terminal regions of host GADs and *Mtb*GadB underscores its potential as a selective therapeutic target (Supplementary Figure S11). Collectively, these findings provide compelling experimental evidence that the CTD of *Mtb*GadB represents a strategically exploitable vulnerability, offering a robust framework for the rational design of specific inhibitors that impair pathogen survival without affecting host homologs.

**Figure 6:**
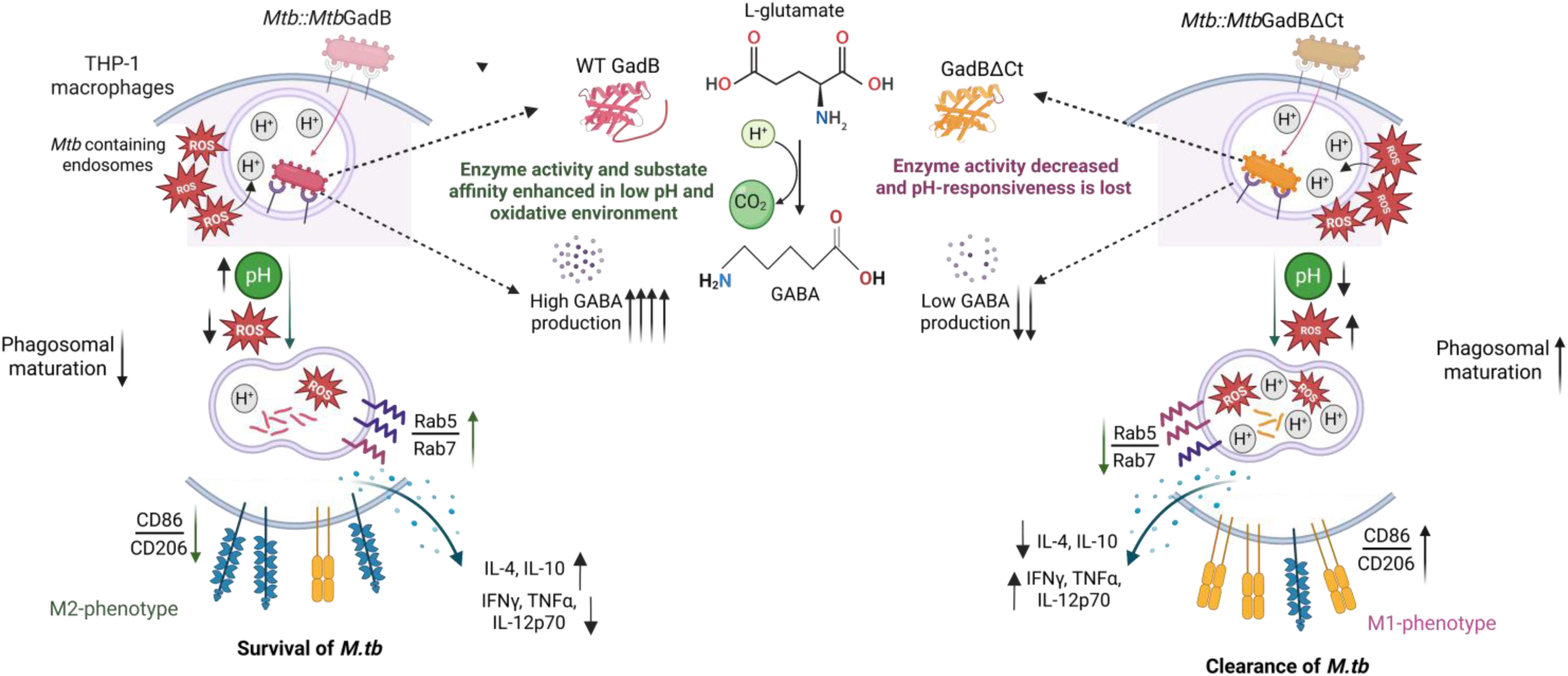
Proposed model of *Mtb*GadB-driven influence of stress-adaptive bacterial metabolic state on macrophage immunomodulation and bacterial persistence: *Mtb*::*Mtb*GadB shows enhanced catalytic efficiency under acidic and oxidative environments of the *Mtb*-containing endosomes, resulting in increased production of GABA. Infection of THP-1 macrophages with the high-GABA- producing (*Mtb*::*Mtb*GadB) strain shows neutralization of pH, and a decrease in cellular ROS, with a high Rab5/7 ratio, leading to a decrease in phagosomal maturation and enhanced bacterial adaptation to these stresses. In parallel, increased *Mtb*GadB activity and the associated rise in intracellular GABA levels drive macrophages toward an anti-inflammatory state, characterized by reduced secretion of pro-inflammatory cytokines (TNF-α, IFN-γ, and IL-12p70) and increased expression of the M2-associated surface marker (CD206), indicative of an M2-like phenotype. Collectively, the dual role of *Mtb*GadB in intracellular stress adaptation and induction of M2-like polarisation promotes enhanced persistence of *Mtb* within macrophages. In contrast, *Mtb*::*Mtb*GadBΔCt shows decreased catalytic efficiency and no pH responsiveness, resulting in less GABA production. Infection with this mutant strain resulted in low pH and accumulation of cellular ROS, along with a low Rab5/7 ratio, leading to an increase in phagosomal maturation. *Mtb*::*Mtb*GadBΔCt strain results in a pro-inflammatory response associated with enhanced expression of M1-associated surface marker (CD86), and decreased anti-inflammatory cytokines (IL-4, IL-10). Together, the inability to counter intracellular acidic and oxidative stress, along with a pro-inflammatory immune response, leads to enhanced clearance of *Mtb* within macrophages.

## Materials and methods Cell lines and antibodies

THP-1 monocytes (ATCC) were maintained in complete RPMI 1640 (Gibco, USA) media, supplemented with 10% Fetal Bovine Serum (Gibco, USA), 1x concentration of Antibiotic-Antimycotic (Gibco, USA) at 37°C in 5% CO_2_ incubator (Eppendorf, Germany). Different antibodies used in this study include anti-GroEL2 (BEI resources); anti-*Mtb*GadB antibody (in-house generated); anti-Rab5 (#3547S Cell Signaling Technology), anti-Rab7 (#9367S Cell Signaling Technology) and anti-GAPDH (#sc-47724 Santa Cruz Biotechnology); Anti-His (#sc-8036 Santa Cruz Biotechnology), Anti-mCherry (#43590S Cell Signaling Technology). Secondary antibodies used in the study include anti-rabbit (#111-035-144, Jackson ImmunoResearch, UK) and anti-mouse (#115-035-146, Jackson ImmunoResearch, UK). All antibodies were diluted in 1x TBST buffer before use.

## Plasmid constructs and mutants

*Mtb*GadB ORF was PCR amplified using H37Rv genomic DNA as template and cloned into the pET28a vector (SnapGene), pMSP12 vector using EcoRI/HindIII (NEB, USA), and pVV16 vector using NdeI/HindIII (NEB, USA) restriction sites to generate pET28a_*Mtb*GadB, pMSP12_*Mtb*GadB and pVV16_*Mtb*GadB, respectively. A catalytic site mutant (Histidine276-Lysine277 mutated to Alanine) was generated by site-directed mutagenesis using the Q5 DNA polymerase (NEB, USA) enzyme, following the manufacturer’s protocol. A C-terminal deletion mutant was generated by cloning truncated *Mtb*GadB into pET28a and pVV16/pMSP vector using specific primers (Table S1). All constructs were confirmed by double digestion and sequencing.

## Bacterial strains

*Mtb::pVV16, Mtb::Mtb*GadB, *Mtb::Mtb*GadBSDM, and *Mtb::Mtb*GadBΔCt strains were generated by electroporation (Eppendorf, Germany) of pVV16 vector, *Mtb*GadB_pVV16, *Mtb*GadBSDM_pVV16, and *Mtb*GadBΔCt_pVV16 into electrocompetent H37Rv cells at a pulse of 2500mV for 4.5 milliseconds, followed by recovery in 7H9 media and plating on 7H10 hygromycin (Himedia, India) plates. Colonies were picked and inoculated in 7H9 media, and expression was confirmed by western blotting using specific antibodies. Similarly, *Msmeg*::pMSP, *Msmeg::Mtb*GadB, *Msmeg::Mtb*GadBSDM, and *Msmeg::Mtb*GadBΔCt strains were generated. All the strains used in the study are tabulated in Table S2.

## Mycobacterial culture and stress treatment

*Mycobacterium tuberculosis H37Rv* or *Mycobacterium smegmatis* was cultured in 7H9 broth (Himedia, INDIA) including 0.4% (v/v) glycerol (Himedia, India) and 0.05% (v/v) Tween 80 (Sigma Aldrich, USA) supplemented with 10% Middlebrook OADC (Himedia, India) at 37°C, 180 rpm until log phase, to an O.D. of 0.7-0.8 at 600nm. Bacterial cells were further harvested, washed with 1x PBS, and resuspended in the required stress media for 36 hours (*Mtb H37Rv*) or 4 hours (*Msmegmatis*). The stress media were prepared mimicking the infection stress conditions, such as acidic stress (pH 5.5), oxidative stress (10 mM H_2_O_2_), in Sauton’s minimal medium (18). Any possible contamination in the culture was ruled out by using the Zeil-Neilson staining kit (Himedia, India) before and after stress. Each experiment was performed in triplicate with technical duplicates. After 36 hours of stress, the cells were harvested for further experiments.

## Homology modeling of *Mtb*GadB

Homology modeling of *MtbGadB* was performed using the Prime application in Schrodinger Maestro Suite 2024 (Schrodinger, LLC, New York, NY, 2024). The 460-amino-acid sequence of *Mtb*GadB (accession ID: I6YG46) was retrieved from the UniProt database. Using NCBI Protein BLAST against the PDB database, *E.coli* GadB protein was identified as the best template, based on high sequence identity and similarity (44.1% and 60.5%) and conserved catalytic residues. Thus, the 3-D crystal structure of the *E.coli* GadB protein (PDB ID: 1pmm; resolution 2.0 Å) was retrieved from the RCSB PDB database, and homology modeling was done using the knowledge-based model-building method in Prime, generating one model. Stereochemical quality was evaluated with Ramachandran plots and structural validation tools. The resulting model displayed a well-preserved fold characteristic of PLP-dependent decarboxylases, with the active-site pocket and cofactor-binding region closely resembling those of the template enzyme. The generated model was selected for further refinement, comprising loop refinement and energy minimization using the OPLS4 force field in Schrodinger Prime. This model was optimized prior to refinement using the protein preparation workflow in Schrodinger Maestro Suite 2024.

### *In silico* alanine scanning

To validate the importance of the selected single-point mutation, *in silico* alanine scanning was performed using the BioLuminate module in the Schrödinger Suite. For this purpose, the complex of *Mtb*GadB with PLP cofactor was considered. In this step, the backbone atoms were optimized, and rotamer prediction was performed for the residues surrounding the site of substitution. The effect of each substitution mutation on the overall stability of the complex was evaluated after local relaxation of the residues around the substituted site. The difference in the binding free energy between the wild-type (ΔG*_wild-type_*) protein and its alanine mutant (ΔG*_alanine_*) counterpart, ΔΔG*_binding_* (kcal/mol), is given by Eq 1:

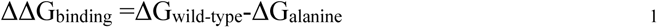

## Mycobacterial infection and persistence assay

THP-1 monocytes (ATCC) were seeded at an appropriate density and differentiated into macrophages using PMA (Sigma-Aldrich, USA) at 100 ng/mL for 24 hours, followed by 12 hours of incubation in 5% CO_2_ at 37°C. Differentiated cells were infected with appropriate mycobacterial strains at MOIs of 1:10 (*Mtb H37Rv*) and 1:20 (*Msmeg*) for 4 hours, followed by stringent washes with 1x PBS to remove the extracellular bacilli. The cells were incubated at 37°C in complete RPMI and harvested at 24 hours post-infection (hpi) (*Mtb H37Rv*), for lysate preparation and RNA isolation. For mycobacterial persistence assay, the infected THP-1 macrophages were lysed (at 0hpi and 24hpi) by incubation in Milli-Q water for 1 hour at 37°C, followed by plating on 7H10 hygromycin agar plates for CFU count. The obtained colonies were counted and the CFU was calculated using the formula cfu/ml= no of colonies*dilution factor/volume of the culture plated. Data was normalised to control infection (*Mtb*::pVV16) and represented as percentage survival.

## Macrophage polarisation

THP-1 monocytes (ATCC) were seeded at an appropriate density and differentiated into macrophages using PMA (Sigma-Aldrich, USA) at 100 ng/mL for 24 hours, followed by 12 hours of incubation in 5% CO_2_ at 37°C. For M1 polarisation, cells were treated with 1μg/ml LPS (#sc3535A) and 20ng/ml IFN-γ (BD #285-IF/CF) for 24 hours. For M2 polarisation, cells were treated with 40ng/ml of IL-4 (BD #204-IL/CF) for 24 hours. After 24 hours of treatment, cells were washed thrice with pre-warmed 1x PBS and incubated at 37°C, 5% CO2 in complete RPMI for 24 hours (44).

## RNA isolation and qRT-PCR

*Mtb H37Rv* cells were harvested after 36 hours of stress and resuspended in Trizol (Invitrogen, USA) along with 0.1mm of zirconia beads. The cells were lysed by bead beating for 15 cycles with a pulse of 1 minutes followed by incubation on ice for 2 minutes. The clear lysate was collected and total RNA was isolated following the manufacturer’s protocol. For THP-1 macrophages, cells were washed with 1x PBS and total RNA was isolated using Trizol method. 5 µg of isolated RNA was treated with DNase I (NEB, USA) to remove any genomic DNA contamination. 1ug of DNase I (NEB, USA) treated RNA was converted to cDNA using the iScript cDNA synthesis kit (BIO-Rad, USA) according to the manufacturer’s protocol. The diluted cDNA was used as template and qRT-PCR was performed using the iTaq Universal SYBR green Supermix (Bio-Rad, USA) following the manufacturer’s protocol, in a 96-well plate (Applied Biosystems, USA) and analyzed using QuantStudio3 software.

## Western blotting

The cells were resuspended in 1x PBS (mycobacterial) or RIPA buffer (mammalian) and lysed by bead beating (mycobacterial) or vigorous vortexing (mammalian). The clear lysate was collected and protein concentration was estimated by Bradford assay (Bio-Rad, USA). Equal amounts of lysates were loaded on SDS-PAGE, transferred onto a nitrocellulose membrane (0.2 μm Nitrocellulose membranes, Bio-Rad, USA) and visualized by ponceau (Fluka Analytical, USA) staining. The blots were blocked with 5% skimmed milk (Himedia, India) for 1 hour at room temperature, followed by 3 washes with 1x PBS and then incubated with primary antibody overnight at 4°C. The following day, the blots were washed 3 times with 1x TBST, incubated with a secondary antibody (Jackson ImmunoResearch, UK) for 2 hours at room temperature, and developed using a WesternBright™ ECL kit (Advansta, K-12045-D20) on a Bio-Rad ChemiDoc instrument. The protein levels of *Mtb*GadB were quantified by densitometry analysis using ImageJ software and normalized to the loading control for each condition.

## Metabolite isolation

*Mtb H37Rv* cells were harvested after 36 hours of stress, and metabolism was quenched by freezing in liquid nitrogen. Culture supernatant was collected and passed through 0.22μm filters. For intracellular metabolite isolation, the cells were thawed on ice for 5 minutes and resuspended in metabolite isolation solution (2:1 methanol: chloroform). The cells were lysed by bead beating for 15 cycles with a 1-minute pulse, followed by incubation on ice for 2 minutes on a mini bead-beater. The clear lysate was collected by high-speed centrifugation. For extracellular metabolites, 3-4 volumes of chilled methanol were added to the culture filtrate, followed by mild vortexing and incubation at -80°C for 2 hours. The sample was further centrifuged at 14,000xg for 15 minutes at 4°C, and the supernatant was collected. All samples were vacuum-concentrated using a SpeedVac concentrator (Eppendorf, Germany) and stored at -80°C until quantification or acquisition. For THP-1 macrophages, the cells were washed with pre-warmed 1x PBS, and 1 ml of chilled metabolite quenching solution (50% methanol: 30% acetonitrile: 20% water) was added per 1 million cells, and the mixture was incubated on ice for 30 minutes. Cells were collected using a mechanical scrapper (#3010 Corning, USA) and vortexed at 4°C for 15 minutes, followed by centrifugation at high speed (>13,000rpm) for 15 minutes at 4°C. Extracted metabolites were vacuum concentrated using Speed Vac concentrator (Eppendorf, Germany) and stored at -80°C until quantification or acquisition.

## NMR spectroscopy to measure metabolites

Dried bacterial metabolites were dissolved in 550 μL phosphate buffer (H_2_O: D_2_O, 20:80 v/v) containing sodium trimethylsilyl propionate (TSP, 0.25 mM) and transferred to NMR tubes. ^1^H NMR spectroscopy was performed on a 600 MHz spectrometer (Bruker Biospin, Ettlingen, Germany) using a triple-resonance probe equipped with a Z-axis gradient. Spectra were acquired using a pulse-acquire sequence with the following parameters: 32,768 data points, spectral width of 7212 Hz, repetition time of 6 sec, and 32 scans. Water suppression was achieved by pre-saturation during the relaxation delay (4 sec). Free induction decays (FIDs) were apodized using a Lorentzian window function (line broadening = 0.5 Hz), zero-filled to 262,144 points, and Fourier transformed, followed by phase and baseline correction. Metabolite intensities were quantified relative to TSP, and concentrations were calculated using TSP (0.25 mM) as an internal standard.

## Protein purification and antibody generation

*E.coli* BL21-pET28a_*Mtb*GadB, *E.coli* BL21-pET28a_*Mtb*GadBSDM, and *E.coli* BL21-pET28a_*Mtb*GadBΔCt were cultured in LB broth (Himedia, India) containing 34 µg/mL kanamycin (Himedia, India), at 37°C with shaking at 180 rpm until the optical density at 600 nm (OD_600_) reached 0.6–0.8. Protein expression was induced by adding IPTG (Sigma Aldrich, USA) to a final concentration of 0.2 mM, followed by incubation at 18°C overnight. The bacterial culture was then harvested, and the resulting pellet was resuspended in lysis buffer containing 50 mM Tris (pH 8.0) and 1 mM PMSF (Sigma Aldrich, USA). The cells were lysed using an ultrasonic disruptor under ice bath conditions (34% amplitude, 10 sec pulses ON, 20 sec pulse OFF for 20 minutes at 4°C), followed by centrifugation at 4°C. *Mtb*GadB was purified using Ni-NTA affinity chromatography. The clear lysate was collected and incubated with Ni-NTA beads (Qiagen, Germany) on an endo-rotor at 4°C for 2 hours. Beads were washed with 50 mM Tris (pH 8), 1 mM PMSF, 300 mM NaCl, and 20 mM imidazole, and the protein was eluted with a 250 mM imidazole solution. The protein was dialyzed to remove excess salts and analyzed by 12% SDS-PAGE. Protein concentration was determined by the Bradford protein assay (Bio-Rad, USA) using the manufacturer’s protocol.

Anti-*Mtb*GadB antibody was raised as per the protocol approved by the Institutional Animal Ethics Committee (UH/IAEC/SB/2022/45). Briefly, 500 μg purified *Mtb*GadB was diluted in a mixture of 0.5 ml PBS and 0.5 ml complete Freund’s adjuvant and injected subcutaneously into a healthy rabbit as a prime dose. A booster dose was given every 2 weeks for 3 months, with a gradual decrease in the amount of *Mtb*GadB protein, emulsified with incomplete Freund’s adjuvant to elicit a robust immune response. Once the immunization schedule was completed, approximately 10 ml of blood was collected, and serum was isolated by centrifugation at 2000 x g for 10 minutes. The specificity of the antibody was confirmed by western blotting against purified *Mtb*GadB protein, with a non-specific protein as a negative control (Supplementary Figure S10).

## Enzymatic reaction and GABA quantification (modified Berthelot method)

The activity of *Mtb*GadB was estimated based on the amount of GABA produced by a modified Berthelot reaction (32, 45–47). The reactions were performed under three different conditions: control (pH 7.2), acidic (pH 5.5), and oxidative (0.5 μM H_2_O_2_). Briefly, the 400 μl reaction mixture composed of 0.2 M Na_2_HPO_4_-citric acid buffer (pH was adjusted to 5.5 and 7.2), 12.5 mM Pyridoxal-5’ phosphate (Sigma Aldrich, USA), 50 μg purified protein, and different concentrations of substrate, L-Glutamate (0 mM-100 mM) (Sigma Aldrich, USA), volume of 400 μl is adjusted with MilliQ water. The reaction mixture was incubated at 37°C for 60 minutes, followed by boiling for 10 minutes to inactivate the enzyme. Immediately after boiling, the mixture was cooled on ice. The prepared reaction mixtures were then used to quantify GABA using the Berthelot method. Specifically, 75 μl of the reaction mixture was transferred into a test tube, followed by the addition of 50 μl of 0.2 M borate buffer (pH 9.0), 250 μl of 6% phenol, and 200 μl of 5% sodium hypochlorite (Himedia, India). The contents were thoroughly mixed, heated and rapidly cooled in an ice bath for 10 minutes. Finally, 500 μl of 60% ethanol was added to the reaction mixture, and the absorbance was measured at 645 nm using a UV-Vis spectrophotometer.

The total amount of GABA obtained in the enzymatic assay was measured using GABA (Sigma-Aldrich, USA) as a standard and expressed as micromolar GABA produced per minute. To obtain the kinetic parameters *V*_max_, *K*_m_, and *k*_cat_ of *Mtb*GadB activity, the initial velocities of GABA release assayed at different substrate concentrations were analyzed through Michaelis-Menten plots.

## High Performance Liquid Chromatography

Amount of GABA in a sample was quantified by Reverse-phase HPLC using a Phenomenex Luna 5u C18(2) 100A column (250 × 4.6 mm) as the stationary phase and sodium acetate buffer (pH 9.0) as the mobile phase (48). Briefly, the sample or the enzymatic reaction mixture was mixed with HPLC-grade water and derivatized with the orthophthaldehyde-N-Acetyl Cystine (OPA-NAC) derivatization reagent. The sample was incubated on ice for 2 minutes, and the desired volume was made with HPLC-grade water. The sample was run on a C18 column, with 100% Sodium acetate buffer as mobile phase at room temperature. The chromatogram was recorded at a wavelength of 334nm. GABA (Sigma-Aldrich, USA) and Glutamate (Sigma-Aldrich, USA) were used as standards.

## Fluorescence spectroscopy

To measure the binding between the ligand (Glutamate) and the protein *Mtb*GadB at different pH levels, we used previously validated method (49). The reaction was set up with recombinant Glutamate Decarboxylase at a concentration of 1 µM along with L-glutamate within a concentration range of 0 mM to 50 mM, keeping PLP concentrations constant in the reaction. The final volume of each reaction was adjusted with 0.2 M Na2HPO_4_-citric acid buffer at pH 5.5 and 7.2. Control reactions were performed with MilliQ water and PBS. Intrinsic fluorophore residues were excited at a wavelength of 285 nm, and emission spectra were collected from 300 nm to 420 nm using a Spectrofluorometer FP-8550 (Japan). Spectral readings were monitored and collected, and the graphs were plotted in GraphPad Prism.

## Mycobacterial growth assay

*Mycobacterium tuberculosis H37Rv* was cultured as described earlier. Briefly, the bacteria were grown in 7H9 broth including 0.4% (v/v) glycerol and 0.05% (v/v) Tween 80, supplemented with 10% Middlebrook OADC, at 37°C and 180 rpm until log phase, to an O.D._600_ of 0.7. Cells were then collected, washed with 1x PBS, and equally resuspended in respective stress media. The bacterial cultures were then placed in an incubator at 37°C and 180 rpm, and the O.D._600_ was measured at different intervals up to the specified time.

## Bacterial ROS measurement

The assay was performed with non-pathogenic *Msmeg* strains overexpressing *Mtb*GadB, *Mtb*GadBSDM and *Mtb*GadBΔCt cloned in pMSP shuttle vector. Mycobacterial strains were grown in complete 7H9 media up to log phase and subjected to oxidative stress (10 mM H_2_O_2_) for 4 hours. Total intrinsic ROS was quantified using CM-H_2_DCFDA dye (#C6827 Invitrogen, USA) according to the manufacturer’s protocol. Briefly, mycobacterial strains were grown till log phase and subjected to oxidative stress. After 4 hours of stress, the bacterial cells were collected and washed thrice with pre-warmed 1x PBS, and incubated with CM-H_2_DCFDA (Sigma Aldrich, USA) dye for 30 minutes at 37°C, 180 rpm. The cells were then collected and washed with 1x PBS to remove the excess dye and resuspended in 1x PBS to measure the intrinsic fluorescence. Total fluorescence was recorded at 527 nm in a UV-Vis Fluorescence Spectrophotometer (Eppendorf, Germany) and a graph was plotted using GraphPad Prism.

## Live cell imaging

The assay was performed with non-pathogenic *Msmeg* strains overexpressing *Mtb*GadB, *Mtb*GadBSDM and *Mtb*GadBΔCt cloned in pMSP shuttle vector (cherry tag). THP-1 monocytes were seeded in live-cell imaging dishes (Ibidi GmbH, Germany) at an appropriate density and differentiated into macrophages with PMA at 100 ng/ml for 24 hours, followed by 12 hours of incubation at 37°C with 5% CO_2_. Cells were infected with mycobacterial strains (*Msmegmatis*) at an MOI of 1:20 for 4 hours, followed by stringent washes with 1x PBS to remove the extracellular bacilli. The cells were incubated at 37°C in complete RPMI media for 6 hours. After 6 hours of infection, the cells were treated with Lysotracker Green DND-26 (#L7526 Invitrogen, USA) at 37°C for 30 minutes following the manufacturer’s protocol. The cells were washed twice with 1x PBS, then replaced with fresh media and subjected to live-cell imaging (Leica confocal microscope). The images were acquired as Z-stacks with an axial step size of 0.3 µm, for 15 minutes at every 30 sec and visualized by Leica Las X software. The images were analysed using Fiji (ImageJ) software, detailed in Supplementary Figure 6 (50, 51).

## Flow cytometry

The assay was performed with non-pathogenic *Msmeg* strains overexpressing *Mtb*GadB, *Mtb*GadBSDM and *Mtb*GadBΔCt cloned in pMSP shuttle vector. THP-1 monocytes were seeded at an appropriate density and differentiated into macrophages using PMA at a concentration of 100 ng/ml for 24 hours, followed by 12 hours of incubation at a 37°C incubator with 5% CO_2_. Cells were infected with mycobacterial strains (*Msmegmatis*) at an MOI of 1:20 for 4 hours, followed by stringent washes with 1x PBS to remove the extracellular bacilli. The cells were then incubated at 37°C in complete RPMI media (containing antibiotics) for 6 hours. After incubation, the cells were washed 3 times with 1x PBS and incubated with CM-H_2_DCFDA (Sigma Aldrich, USA) at 37 °C for 30 minutes in the dark, following the manufacturer’s protocol. The cells were washed, collected using a cell scraper, and passed several times through a blunt 20-gauge needle fitted to a sterile syringe to make single-cell suspensions (without any visible clumps), which were then acquired immediately on a BD Fortessa Flow cytometer to evaluate the total median fluorescence intensity (MFI). Cells treated with 200 μM H_2_O_2_ were used as a positive control for experimental validation. The data were analysed using FACS FlowJo.10 software. Data were normalised to uninfected macrophages and represented as MFI fold change.

## Greiss assay

Nitrite accumulation in culture supernatants was measured as an indicator of nitric oxide (NO) production using the Griess reaction. Briefly, THP-1 monocytes (ATCC) were seeded at an appropriate density and differentiated into macrophages using PMA (Sigma-Aldrich, USA) at 100 ng/mL for 24 hours, followed by 12 hours of incubation in 5% CO_2_ at 37°C. Cells were infected with mycobacterial H37Rv strains at an MOI of 1:10 for 4 hours, followed by stringent washes with 1x PBS to remove the extracellular bacilli. The cells were incubated in 5% CO_2_ at 37°C in complete RPMI for 24 hours. After 24 hours of incubation, culture supernatant was collected and passed through 0.22 μm filters and used to measure the total nitrite produced. Equal volumes (50 µL) of culture supernatant and Griess reagent [1% sulfanilamide in 5% phosphoric acid and 0.1% N-(1-naphthyl) ethylenediamine dihydrochloride (NEDA)] were mixed in a 96-well flat-bottom plate and incubated at room temperature for 15 minutes in the dark. The absorbance was measured at 540 nm using a microplate reader. Nitrite concentrations were determined by comparison with a standard curve generated using serial dilutions of sodium nitrite (NaNO₂) prepared in culture medium.

## Cytokine profiling

THP-1 monocytes (ATCC) were seeded at an appropriate density and differentiated into macrophages using PMA (Sigma-Aldrich, USA) at 100 ng/mL for 24 hours, followed by 12 hours of incubation in 5% CO_2_ at 37°C. Cells were infected with mycobacterial strains at an MOI of 1:10 for 4 hours, followed by stringent washes with 1x PBS to remove the extracellular bacilli. The cells were incubated in 5% CO_2_ at 37°C in complete RPMI for 24 hours. After 24 hours of incubation, culture supernatant was collected and passed through 0.22 μm filters and used to measure the different cytokines by indirect ELISA using human BD OptEIA ELISA sets (#555194, #555157, #555183, #555212, #555142) using the manufacturer’s protocol. M1 and M2 polarised macrophages were used as positive controls.

## Cell surface marker staining

THP-1 macrophages were infected with mycobacterial strains as described above and stained with cell surface marker antibodies using the manufacturer’s protocol. Briefly, infected/treated macrophages were washed with pre-warmed 1x PBS and harvested using a mechanical scrapper followed by centrifugation at 1000 rpm for 5 minutes. Non-specific binding was restricted by blocking the cells with Fc blocker (BD Pharmingen #564219) at room temperature for 20 minutes. Cells were washed with 1x PBS and stained with FITC-tagged anti-CD86 (BD Pharmingen #555657) and PE tagged anti-CD206 (BD Pharmingen #555954) for 1 hour on ice in the dark. The cells were washed and fixed using 4% paraformaldehyde (Sigma Aldrich, USA) for 15 minutes in the dark. The cells were washed and passed several times through a blunt 20-gauge needle fitted to a sterile syringe to make single-cell suspensions (without any visible clumps), which were then acquired immediately on a BD Fortessa Flow cytometer to evaluate surface staining. M1/M2 polarised cells were used as positive controls.

## Statistical significance

All the experiments were performed in biological triplicate. Plots were generated, and the statistical significance was calculated by a two-tailed unpaired Student’s t-test using GraphPad Prism. A p-value of =/<0.05 was considered statistically significant.

## Supplementary Information

Supplementary figures S1-S11 Supplementary tables S1-S4

### Funding acquisition

Science and Research Engineering Board, India (SERB) Research grant (SPR/2021/000137) under the scheme SUPRA, and the DST-BRICS grant DST/ICDBRICS/PilotCall2/MRP-TB (G) to SB.

## Acknowledgments

We acknowledge DST-INSPIRE JRF and SRF to AA. SS acknowledges support from DST/WISE-PDF/CS-22/2023. We thank the Department of Biochemistry and the School of Life Sciences for their common facilities. We thank the UoH-NIAB BSL3/ABSL3 facility for aiding mycobacterial work. We acknowledge Novelgene Technologies for the LC-MS acquisition. We acknowledge the support from BEI resources for the GroEL2 antibody. We thank Mr. Vinay for helping with HPLC. We thank Mr. Manikanta for assisting with flow cytometry. We thank Ms. Deepti for helping with microscopy. We thank Mr. Omkar Zade for assisting in antibody generation. We thank Mr. Asutosh Sahoo for helping in protein purification. DST-FIST to the Department of Biochemistry (SR/FST/LS-II/2023/1172), the DBT-BUILDER grant to the School of Life Sciences (BT/INF/22/SP41176/2020), and the Institution of Eminence supported projects RC1-20-017 and RC4-21-012 to SB and IoE to the University of Hyderabad MHRD (F11/9/2019-U3(A) for infrastructure support to the department of Biochemistry, School of Life Sciences and University common facilities are acknowledged. The model figure was generated using the BioRender.com site. We acknowledge GraphPad Prism for aiding the generation of graphs and statistical analysis, and AI tools, Grammarly for language review and correction.

## Author Contributions

AA: conceptualization, investigation, designed and performed experiments, analysed data, manuscript writing (original draft), review and editing; AM: performed experiments, analysed data, manuscript review and editing; SS: bioinformatics analysis, manuscript review and editing; ABP: NMR acquisition and analysis, manuscript review and editing; KM: manuscript review and editing; SB: conceptualization, supervision, manuscript review and editing, funding acquisition.

## Conflicts of Interest

The authors declare no conflict of interest.

